# 4C-ker: A method to reproducibly identify genome-wide interactions captured by 4C-Seq experiments

**DOI:** 10.1101/030569

**Authors:** Ramya Raviram, Pedro P. Rocha, Christian L. Müller, Emily R. Miraldi, Sana Badri, Yi Fu, Emily Swanzey, Charlotte Proudhon, Valentina Snetkova, Richard Bonneau, Jane Skok

## Abstract

4C-Seq has proven to be a powerful technique to identify genome-wide interactions with a single locus of interest (or “bait”) that can be important for gene regulation. However, analysis of 4C-Seq data is complicated by the many biases inherent to the technique. An important consideration when dealing with 4C-Seq data is the differences in resolution of signal across the genome that result from differences in 3D distance separation from the bait. This leads to the highest signal in the region immediately surrounding the bait and increasingly lower signals in far-*cis* and *trans*. Another important aspect of 4C-Seq experiments is the resolution, which is greatly influenced by the choice of restriction enzyme and the frequency at which it can cut the genome. Thus, it is important that a 4C-Seq analysis method is flexible enough to analyze data generated using different enzymes and to identify interactions across the entire genome. Current methods for 4C-Seq analysis only identify interactions in regions near the bait or in regions located in far-*cis* and *trans*, but no method comprehensively analyzes 4C signals of different length scales. In addition, some methods also fail in experiments where chromatin fragments are generated using frequent cutter restriction enzymes. Here, we describe 4C-ker, a Hidden-Markov Model based pipeline that identifies regions throughout the genome that interact with the 4C bait locus. In addition we incorporate methods for the identification of differential interactions in multiple 4C-seq datasets collected from different genotypes or experimental conditions. Adaptive window sizes are used to correct for differences in signal coverage in near-bait regions, far-*cis* and *trans* chromosomes. Using several datasets, we demonstrate that 4C-ker outperforms all existing 4C-Seq pipelines in its ability to reproducibly identify interaction domains at all genomic ranges with different resolution enzymes.

**AUTHORS SUMMARY:** Circularized chromosome conformation capture, or 4C-Seq is a technique developed to identify regions of the genome that are in close spatial proximity to a single locus of interest (‘bait’). This technique is used to detect regulatory interactions between promoters and enhancers and to characterize the nuclear environment of different regions within and across different cell types. So far, existing methods for 4C-Seq data analysis do not comprehensively identify interactions across the entire genome due to biases in the technique that are related to the decrease in 4C signal that results from increased 3D distance from the bait. To compensate for these weaknesses in existing methods we developed 4C-ker, a method that explicitly models these biases to improve the analysis of 4C-Seq to better understand the genome wide interaction profile of an individual locus.

## Introduction

Understanding the 3D organization of the genome and the intricacies of chromatin dynamics has been the focus of studies aimed at characterizing gene regulation in physiological processes and disease states [1, 2]. Microscopy based studies provided the first snapshots of nuclear organization, revealing that individual chromosomes occupy distinct territories with little intermingling between them [3, 4]. The development of chromosome conformation capture (3C) transformed the field of nuclear organization enabling identification of chromatin interactions at the molecular level and at the same time opening the door to high-throughput, genome-wide techniques [5]. Hi-C, for example, captures all pairwise interactions in the nucleus and has revealed that chromosomes segregate into two distinct spatial compartments (A and B) depending on their transcriptional and epigenetic status [6]. These compartments are further subdivided into Topological Associated Domains (TADs), which are highly self-interacting megabase scale structures [7-9]. To probe interactions between regulatory elements using Hi-C requires a depth of sequencing that for many labs is cost-prohibitive [10]. 5C can circumvent these issues, but the interaction analysis is limited to the portion of the genome for which primers are designed [11]. Circular chromosome conformation capture combined with massive parallel sequencing (4C-Seq) is currently the best option for obtaining the highest resolution interaction signal for a particular region of interest.

In 4C-Seq, an inverse PCR step allows for the identification of all possible genome wide interactions from a single viewpoint (the “bait”) and an assessment of the frequencies at which these occur. The sequencing coverage obtained by 4C near the bait region is extremely high and therefore enables precise characterization and quantification of regulatory interactions [12, 13]. By focusing on one locus at a time and thus only the interactions that this locus is engaged in, 4C can reproducibly identify long-range interactions on *cis* and *trans* chromosomes [14]. For example, 4C was used to demonstrate that genes controlled by common transcription factors tend to occupy the same nuclear space even when located on different chromosomes [15, 16].

There are many inherent biases specific to the 4C technique that has made detecting meaningful and reproducible interactions challenging. First, in accordance with the chromosome territory model, the majority of 4C signal is located on the bait chromosome. Secondly, coverage and signal strength are highest in the region around the bait and this decreases along the chromosome as a function of linear distance from the bait. Third, the restriction enzyme used for the first digest in the experiment is an important determinant of the resolution at which interactions can be detected. Finally, as with most PCR-based techniques, 4C data includes PCR artifacts that manifest as a large accumulation of reads in particular locations.

Current methods of analysis have addressed some of these issues, however there are still many hurdles to overcome. Specifically, existing methods do not properly account for the differences in 4C signal coverage across the genome and therefore they are only able to either identify interactions in (i) regions where the signal is highest, i.e., near the bait or (ii) regions of low 4C signal (far-*cis* and *trans)*. Thus there is no method that comprehensively identifies interactions across the genome. In addition, most methods were developed and tested using datasets generated with 6bp cutters and we show that they do not perform well with 4bp cutter generated libraries.

The goal of 4C-ker is to address these weaknesses by: 1) identifying domains that interact most frequently with the bait across the genome in a given population of cells, and 2) detecting quantitative differences in 4C-Seq signal between conditions. Here we use a Hidden Markov Model to account for the polymer nature of chromatin, in which adjacent regions share a similar probability of interacting with the bait. In addition, to account for the variation in signal captured at different 3D distances we use a window-based approach. To determine the window size of analysis, we adapted a k-th nearest neighbor approach to account for the decrease in 4C-Seq coverage along *cis* and *trans* chromosomes. We used 4C-ker to analyze several publically available 4C-Seq datasets as well as data generated in our own lab and compared this with other published methods. Our results demonstrate that 4C-ker can correct for multiple 4C-Seq biases and reproducibly detect genome wide interactions from the bait viewpoint. Importantly, 4C-ker is the only tool that can identify interactions with regions in near and far-*cis* as well as *trans*.

## Results

### Workflow of 4C-ker

We developed 4C-ker to identify genome-wide interactions generated by 4C-Seq data and to quantitatively examine differences in interaction frequencies between conditions. The main components of the 4C-ker method are outlined in **Fig 1**. First, 4C-Seq reads are mapped to a reduced genome consisting of unique sequences adjacent to all primary restriction enzyme sites in the genome. Mapping to a reduced genome helps to remove spurious ligation events that do not result from crosslinking. The analysis of 4C-Seq is typically performed separately for *cis* and *trans* chromosomes because of the large differences in signal in these distinct locations. Additionally, we present the option for focusing the analysis on the region surrounding the bait, where 4C-Seq signal and resolution are highest. A window-based approach is applied in order to take into account of differences in near-bait far-*cis* signal strength at different 3D distances and the dynamic nature and variability of chromosome interactions in a population of cells.

**Fig 1:**
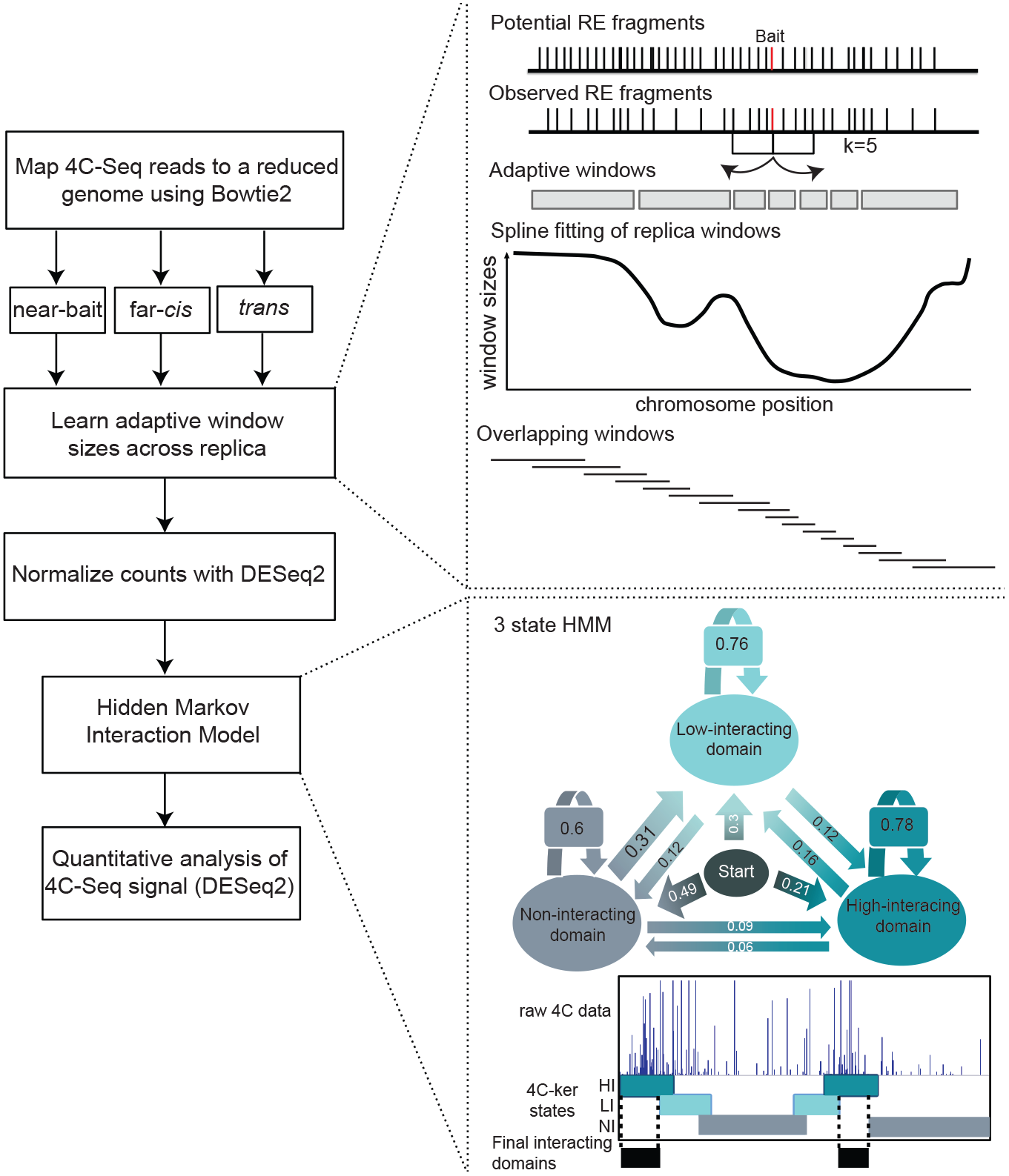
Workflow of 4C-ker. The key features of the method are outlined in the figure. A more detailed explanation of each section can be found in the materials and methods section.

One of the most challenging aspects of 4C-Seq is determining the window size at which the data should be analyzed. Adaptive window sizes that depend on the distance to the bait can adjust for differences in coverage of 4C-Seq signal in regions near the bait, far-*cis* and *trans* chromosomes. 4C signal is generally higher around the bait region and decreases in far-*cis* and *trans*. We developed a kth nearest neighbor method to build overlapping windows of adaptive sizes based on the 4C-Seq signal coverage of a given dataset at each location in the genome. The aim of using this approach is to obtain a similar number of observed fragments for each window. Therefore the size of each window is determined by the amount of signal detected in the region. This will result in small windows near the bait and other regions where there is high coverage, versus larger windows further away from the bait where there is low coverage. An example of this is shown in **S1A Fig**.

To correct for PCR amplification bias, the counts at observed fragments within each window that exceed the 75^th^ quantile for that window are limited to this value. The window counts across all samples are subsequently normalized using DESeq2 [17] to correct for sequencing depth. For the *cis* chromosome, the linear distance from the bait to the mid-point of each window is used to correct for the inverse relationship between counts and linear separation from the bait. The counts and distances are log transformed and are used as inputs (observed states) for the Hidden Markov Model (HMM). A separate model is used for *cis* and *trans* chromosomes (in the latter there is no effect of linear distance from the bait). A three-state HMM is used to partition the genome into windows that interact with the bait at (1) high frequency, (2) low frequency and (3) those that do not interact. Use of overlapping windows, allows us to more precisely define the regions of high interaction (for more detailed explanation of the workflow, refer to the methods section). The resulting parameters for the model show higher probabilities for transitioning to the same state, correctly accounting for the polymeric nature of DNA on chromosomal interactions (**Fig 1**, 3-state HMM).

Since we use overlapping windows, consecutive windows that are detected as highly interacting can be stitched to form a ‘domain’ of high interaction with the bait. Domains that are found as interacting at a high frequency in atleast one sample can be used for downstream quantitative analysis. Furthermore DESeq2 designed for differential analysis of counts-based sequencing data can be used to quantitatively compare interactions across conditions. The 4C-ker pipeline is available as an R package along with the domains of interactions identified for all the datasets analyzed in this study (github.com/rr1859/R.4Cker).

### 4C-ker identifies close range interactions

4C-Seq is commonly used to identify regulatory interactions that occur in close linear proximity to the bait. Therefore, we provide an option to focus the analysis only in this region, where the highest resolution interactions are identified. An important aspect of 4C-Seq library preparation is the choice of restriction enzyme used to digest cross-linked chromatin as the genome-wide frequency of enzyme recognition sites determines the resolution of the experiment. Therefore, 4bp cutters such as DpnII or NlaIII, which cut the genome more frequently, provide a higher resolution profile of 4C interactions compared to 6bp cutters like HindIII (**S1B Fig**) [18]. To ensure that our method works with both types of restriction enzymes, we tested it using several datasets generated from our lab as well as all publically available datasets for which replicates are available that passed our stringent quality controls. See methods section and Supplementary Table 1 for details).

The near-bait analysis was restricted to 10MB around the bait for 6bp cutters and 2MB for 4bp cutters as these are the regions that contain the highest 4C signal in each case. As can be seen in **Fig 2A** and **2B**, raw 4C-Seq signal is highest near the bait and decreases with increasing linear distance. As 4C-ker corrects for this decrease in signal it is able to detect interactions across the entire region analyzed. In addition, due to the adaptive windows, the size of interactions detected are smallest near the bait where coverage is highest and larger in regions further away from the bait where coverage is lower (**Fig 2C** and **2D**). The resolution of domains identified by 4C-ker can be conveniently adjusted by using different values for the number of observed fragments used to generate the adaptive windows (**S2A Fig**). Here we used values ranging from 3-10 in the 2MB region around the bait. As the value of ‘k’ increases, we observe a consistent increase in the size of the domains as well as increased similarity between replicates (see methods section for details). The parameter k can be adjusted by the user depending on the biological question that is being addressed. For example, if the aim of the study is to identify interactions between enhancers and promoters, we suggest k=3-5. In order to identify larger domains that coincide with broad regions encompassing chromatin with similar histone modifications, setting k=10 is a suitable choice.

**Fig 2.**
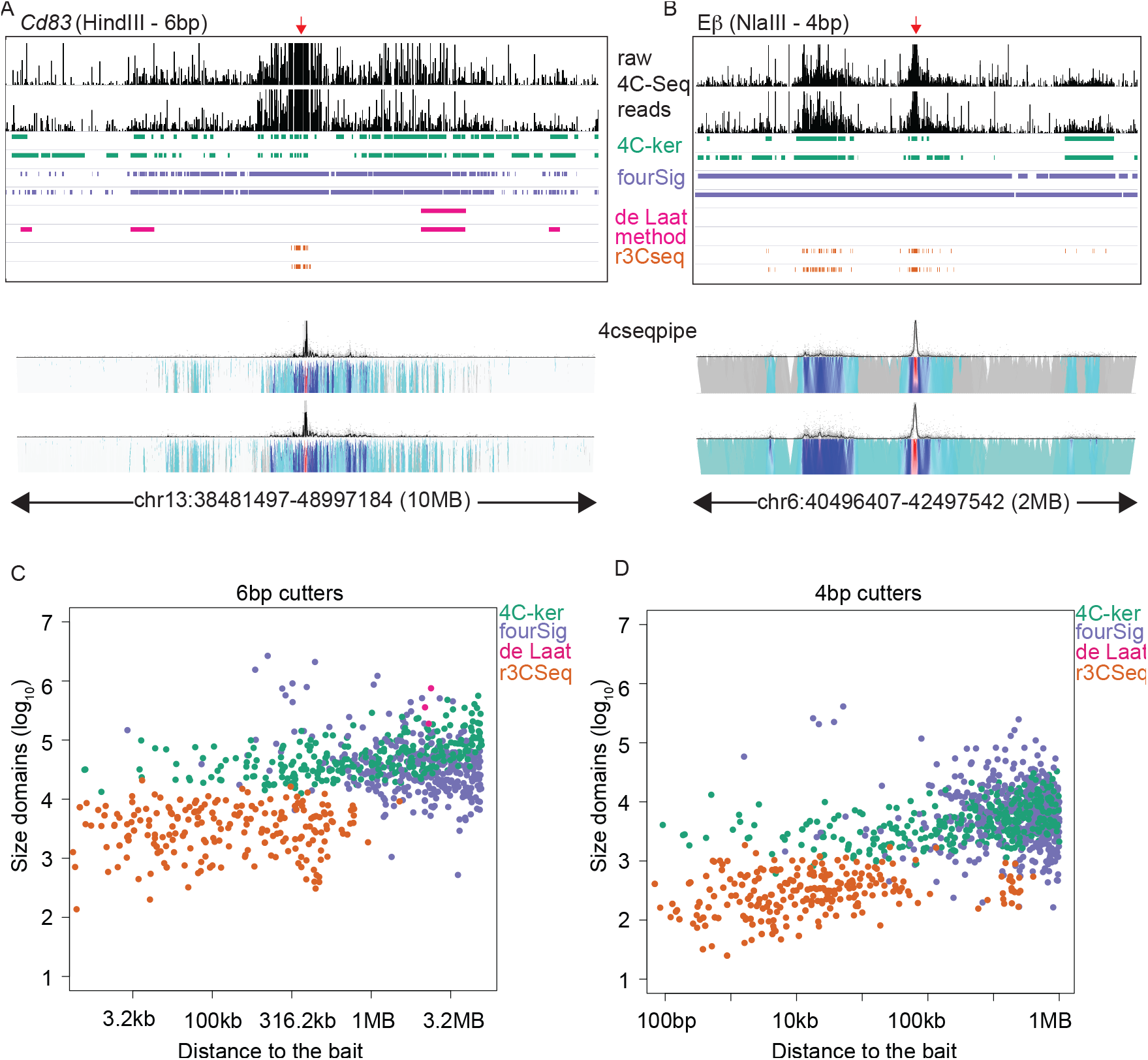
4C-ker outperforms other methods in the region near the bait when the similarity of interacting regions between replicates is examined for four methods. (**A-B**) Example datasets for 6bp and 4bp cutter experiments. Raw 4C-Seq reads are shown for a 10MB region around the bait in (**A**) and 2 MB in (**B**). Experiment in **A** was performed using activated B cells digested with HindIII and a bait near the *Cd83* locus. Experiment in **B** was performed in double negative T cells digested with NlaIII and a bait near the Eβ enhancer of *Tcrb*. Significant interactions determined by each method for 2 replicates are shown below the raw 4C-Seq profile. Domainograms generated using 4cseqpipe are displayed for the same region. (**C-D**) Each dot represents the distance of the midpoint of each interacting domain to the bait plotted against its size. Plots only contain domains that overlap by 50% between replicates.

To assess the performance of 4C-ker we used existing methods to analyze the same datasets. There are currently four publicly available methods to detect significant interactions using 4C-Seq datasets (fourSig, Splinter et al, r3CSeq and FourCSeq). Details of how we implemented these algorithms for comparison with the 4C-ker pipeline can be found in the methods section. Although the method developed by van der Werken et al (4cseqpipe) [13] does not identify significant interactions, it provides a good visualization tool for 4C-Seq signal near the bait (**Fig 2A** and **2B**). The fourSig approach generates windows based on restriction enzyme fragments and compares the counts within each window against a random background distribution [19]. As fourSig does not take account of the impact of distance on 4C-Seq signal, it identifies most of this region as large interacting domains and this results in a high similarity index between replicates (**S2B** **and** **S2C Fig**). However, in contradiction to decreasing resolution of 4C-Seq signal with increased separation from the bait, the size of the domains identified by fourSig are largest near the bait and these decrease with increasing separation from the bait (**Fig 2C** and **2D**). The method described by Splinter et al [20], referred to here as the ‘de Laat method’ excludes the 2MB region around the bait and only calls interactions in the rest of the genome based on enrichment of binary coverage in a given window, compared to a local background. As such, the de Laat method does not identify any interactions with 4bp cutters (**Fig 2B, 2D**). Moreover, using the 6bp cutter datasets it only identifies interactions in 2 out of 7 datasets in the 10MB region (**S2B Fig**). Together these findings reflect the limitations of this method in detecting 4C-Seq interactions in the region with highest coverage, where the majority of important regulatory interactions occur. The r3CSeq method uses reverse cumulative fitted values of the power law normalization and a background scaling method to correct for interactions near the bait [21]. This approach also provides the option to detect interactions at the fragment level or at the window level. In most datasets r3CSeq only identifies significant interaction near the bait as shown in **Fig 2A**, **2C** and **2D** and therefore have a high similarity index between replicates (**S2B** **and** **S2C Fig**). Although interactions further from the bait are identified (**Fig 2B**), they are not reproducible as measured by the similarity index (**S2C Fig**). The FourCSeq pipeline only has the option to analyze interactions at the fragment level. It is based on the DESeq2 method with an additional function that corrects for the effect of linear distance from the bait [22]. This method failed to identify any significant interactions for any of the datasets analyzed. If interactions between regulatory elements are being analyzed, the majority will be identified in the region near the bait. Therefore, it is important that a 4C-Seq analysis can properly identify these interactions. Here we show that 4C-ker outperforms other methods and identifies interactions that correctly reflects the nature of high-resolution 4C-Seq signal in this region.

### 4C-ker identifies long-range interactions

We next used 4C-ker for analysis of the entire bait chromosome using the same fourteen datasets described above. Due to lower 4C-Seq signal in regions distant from the bait (far-*cis*) the correlation between replicates decreases compared to near-bait regions (**S3A Fig**). This difference is more pronounced with 4bp cutter generated datasets. A potential explanation for this difference is that when 4bp cutters are used 4C-Seq coverage in windows distant from the bait decreases at a much faster rate than when using 6bp cutters (**S3B Fig**). Based on these results, it is clear that when designing a 4C-Seq experiment, the biological question should determine the choice of primary restriction enzyme. For example, to detect long-range interactions in *cis* and *trans* it seems preferential to use a 6bp cutter to achieve a more reproducible 4C profile. On the other hand, for characterization of short-range regulatory interactions, 4bp cutters provide a high-resolution map of near-bait interactions, as previously shown [18, 23, 24].

**Fig 3.**
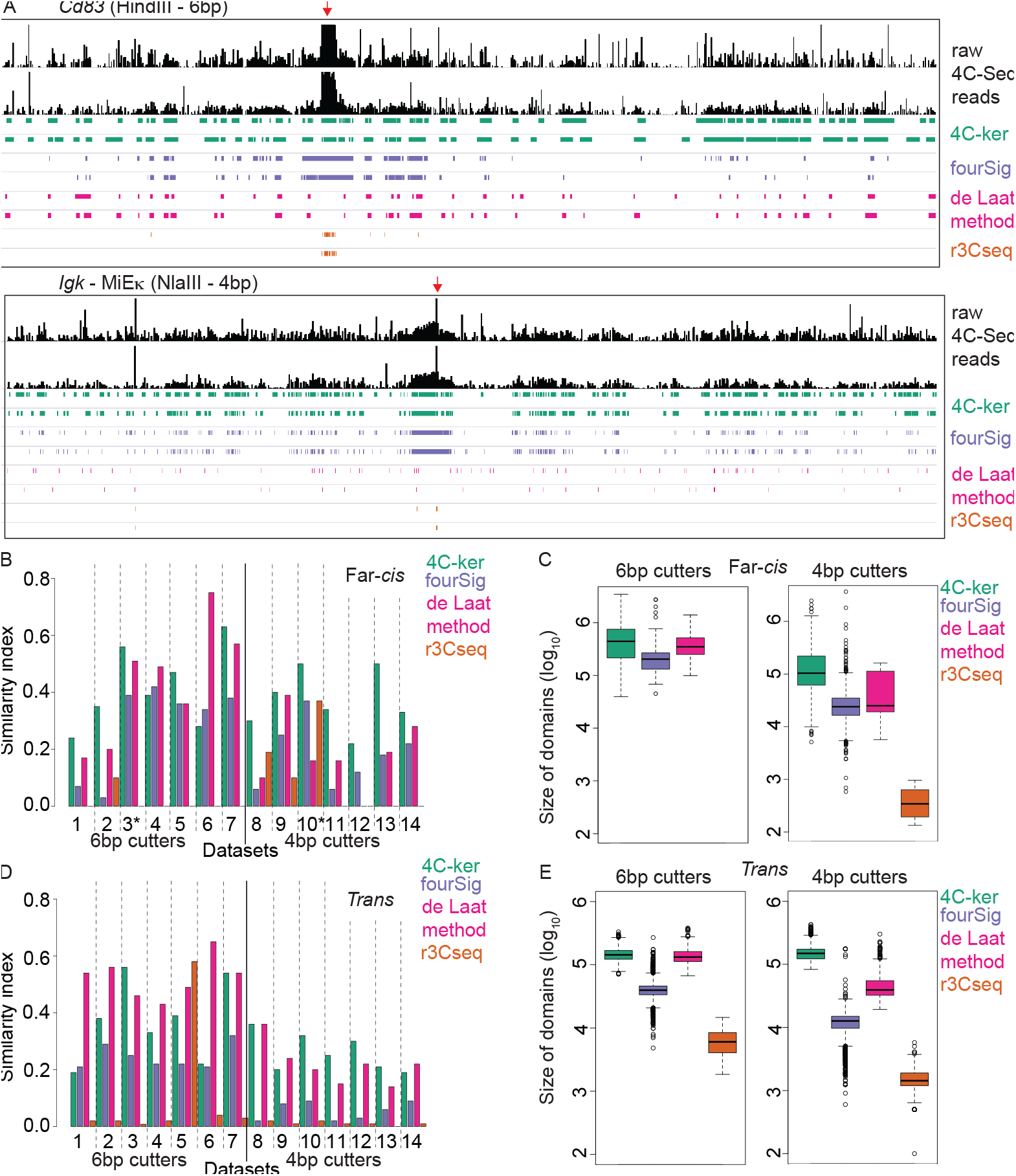
4C-ker identifies the most reproducible interactions across the *cis* chromosome and exhibits stable performance for 4bp and 6bp cutters. (**A**) Example datasets of 6bp and 4bp cutter experiments. Raw 4CSeq reads are shown for the entire bait chromosome. Experiment shown in the top panel in A was performed in activated B cells digested with HindIII using a bait near the *Cd83* gene. Experiment shown in the bottom panel in **A** was performed in immature B cells digested with NlaIII using a bait near the MiEκ enhancer of *Igk*. Significant interactions determined by each method for 2 replicates are shown below the raw 4C-Seq profiles. (**B**) Similarity index between replicates in far-cis. Example datasets shown in **A** are denoted with an asterisk (*) (**C**) Boxplot of the size of domains identified in far-*cis* by each method using all fourteen datasets. (**D**) Similarity index between replicates for domains identified across all *trans* chromosomes. (**E**) Boxplot of the size of all domains in trans identified by the four methods.

With adaptive window sizes and consideration of distance separation from the bait, 4C-ker is able to reproducibly identify domains of interaction across the whole *cis* chromosome. As expected, interacting domains proximal to the bait are smaller in line with the fact that increased 4C-Seq signal allows for generation of smaller windows of analysis. In contrast, in regions located distal to the bait where the 4C-Seq signal is reduced, the window sizes for analysis are increased and 4C-ker identifies larger interacting domains (**Fig 3A**). In comparison to other methods, 4C-ker identifies the most reproducible interactions for all 4 bp cutter datasets and for 5 out of 7 6bp cutter datasets (**Fig 3B**). For 6bp cutter datasets, the domains identified by 4C-ker are comparable in size to the de Laat method (**Fig 3C**). For the 4bp cutter datasets the other methods identify smaller domains compared to 4C-ker, however they do not achieve high similarity indices between replicates. Although the de Laat method and fourSig perform comparably to 4C-ker when analyzing 6bp-generated data they fail to do so with 4bp datasets.

To validate interactions identified by 4C-ker we used the *Igh* Cγ1 HindIII dataset and performed 3D-FISH to analyze interactions with *Igh*. We selected three bacterial artificial chromosome (BAC) probes that hybridize to high, low and non interacting regions in close proximity to each other (4-7Mb), but separated from *Igh* by ∼70Mb (**S4A Fig**). Of note, the selected non interacting region is in closer linear distance to *Igh*, and the highest interacting region is furthest away. Using differentially labeled BAC probes for these regions in conjunction with an *Igh* specific probe we found that in accordance with the 4C-ker output, the BAC in the high interacting domain is in closer spatial proximity to *Igh* than the BACs in the low and non interacting domains (**S4A Fig** and **Fig 3C**). It is of note that all other methods failed to identify the region that we validated by FISH as a significant interaction.

**Fig 4:**
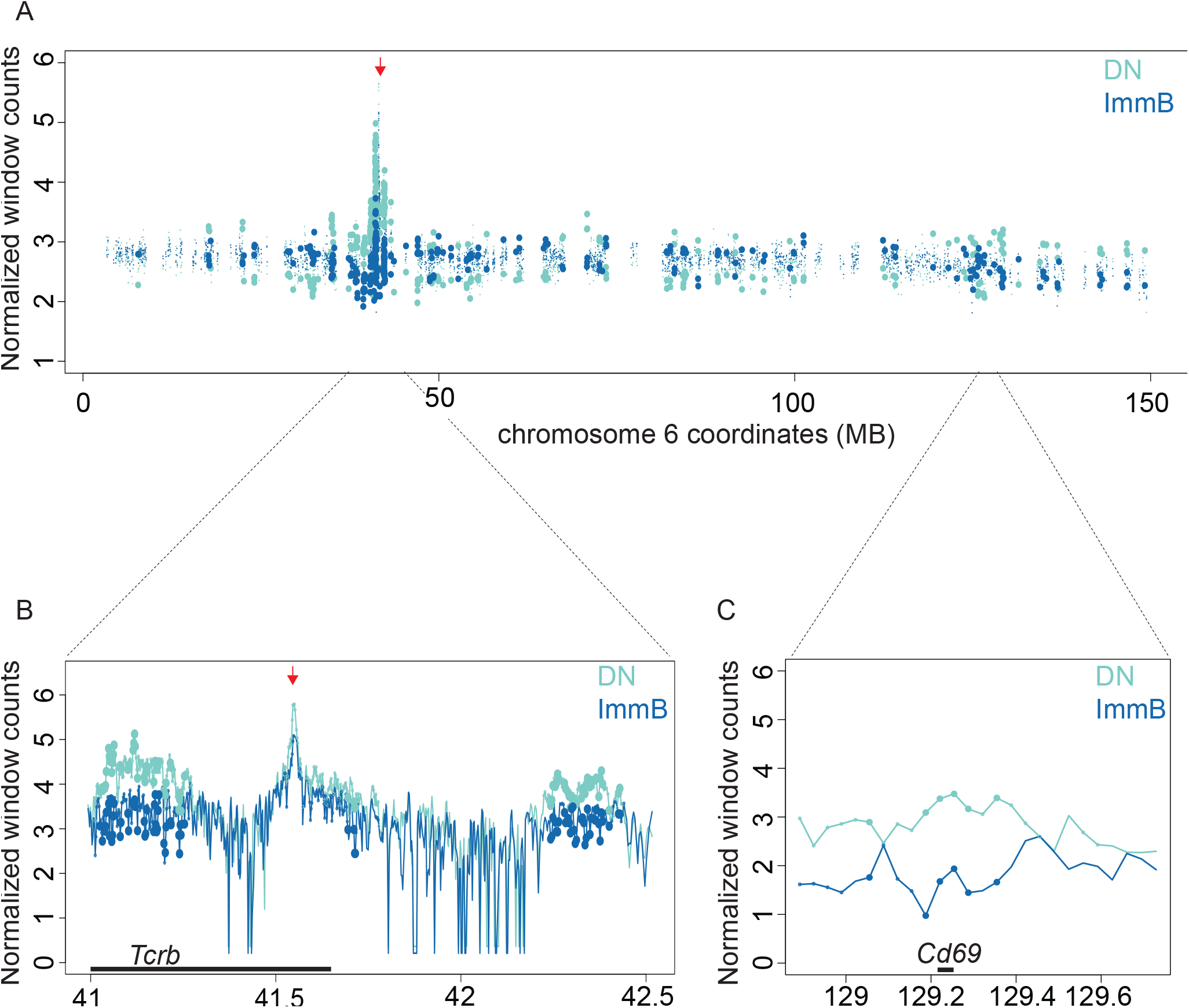
Identification of differentially interacting regions (A-C) Windows located within Dataset-specific Interacting Domains (DIDs) are represented as small circles with the normalized read count for each condition. Large filled circles represent windows with an FDR adjust p-value smaller than 0.05 for the difference between the two cell-types. The red arrow above the plots represents the bait region. Lines represent the average of the replicates for each window across the displayed region. Large filled circles represent windows with an FDR adjusted p-value smaller than 0.05 for the difference between the two cell-types. (A) Whole chromosome, (B) Near the bait, (C) Far *cis* (region around *Cd69*.

According to the chromosome territory model most interactions occur between loci on the same chromosome. As such, inter-chromosomal interactions occur at low frequency. However, unlike other 3C-based techniques, 4C-Seq can still detect these interactions. Nonetheless, since at least 40% of the signal is on the *cis* chromosome, the rest is spread over all *trans* chromosomes and is thus significantly reduced. As a result the 4C signal is less reproducible compared to interactions on the bait chromosome (**S3A Fig**). 4C-ker and the de Laat method outperform fourSig and r3CSeq in identifying *trans* interactions. Both 4C-ker and the de Laat method identify equivalently sized interaction domains across all fourteen (6bp and 4bp cutter) datasets (**Fig 3D** **and** **3E**). In most cases 4C-ker outperforms the de Laat method in identifying reproducible interactions from 4bp cutter experiments, while the reverse is true for most 6bp cutter experiments.

### Differential analysis of interaction profiles across multiple 4C experiments

One useful application of 4C-Seq is a quantitative comparison of interactions from a particular viewpoint across conditions or cell types. The highly interacting domains identified by 4C-ker for several conditions can be merged to generate a list of “Dataset-specific Interacting Domains” (DIDs). These domains represent regions that are interacting with the bait in at least one of the conditions. In general, 4C-Seq counts follow a negative binomial distribution, which is suitable for differential DESeq2 analysis. We use raw counts for the adaptive windows that fall within DIDs and normalize these using DESeq2 to correct for sequencing depth. To identify differential interactions between conditions we use an FDR adjusted p-value of < 0.05.

To test this approach, we compared two datasets generated with NlaIII digested 4C template that have a bait on the Eβ enhancer of *Tcrb* in double negative (DN) and immature B (ImmB) cells. DIDs were generated for the bait chromosome (chromosome 6) for these two cell types. In **Fig 4A**, the normalized values for windows within DIDs are plotted across the entire bait chromosome. Windows that are significantly different in the two cell-types are represented as larger filled circles. It is clear from **Fig 4A** that the majority of differentially interacting regions are concentrated near the bait. This can be seen in detail for the interaction of the Eβ enhancer with the 5’ end of the *Tcrb* gene (**Fig 4B**). In DN cells this locus is in a contracted conformation which brings distal Vβ genes into contact with the proximal DJCβ region for V(D)J recombination [25]. In contrast, the locus does not recombine in B cells and is not in a contracted form and the Vβ genes are found in less frequent contact with the bait. Interestingly, we found a differentially interacting DID in far-*cis* containing the *Cd69* gene (**Fig 4C**), which is a known T cell marker and interacts more frequently with Eβ in DN cells compared to Immature B cells. This is expected since both *Cd69* and *Tcrb* are active in T cells and it has been shown that transcriptionally active regions come into frequent contact [15, 16]. Thus, the DIDs determined by 4C-ker can be used to detect quantitative interactions that correlate with functional processes.

### Long-range interacting regions have similar accessibility and transcriptional profiles to the bait

The ability to detect reproducible long-range interactions with 4C-ker enables us to assess the properties of these regions. Based on nuclear organization principles described by 3C-based studies [6, 15, 16] we validated 4C-ker domains by assessing if they preferentially contact regions with the same transcriptional and epigenetic status as the bait. For this, we used 4C data generated with the Eβ enhancer bait in DN T cells and immature B cells. Using ATAC-Seq [38], a technique that identifies accessible regions of chromatin, we find that as expected, the enhancer is active in T cells and inactive in B cells (**Fig 5A**). Conversely, a bait on the MiEκ enhancer of *Igk* is active in B cells and inactive in T cells (**Fig 5A**). Using 4C-ker we identified the highly interacting domains with each bait across the two cell types. Since we used NlaIII to generate the template we restricted the analysis to the bait chromosome. We then asked if the 4C interacting domains are enriched for ATAC-Seq peaks. Here, we define enrichment as the ratio of the sum of the size of ATAC-Seq peaks within interacting regions to those within a background generated by randomly repositioning these domains along the chromosome. In T cells, where the Eβ enhancer is active, we found a higher enrichment of ATAC-Seq peaks in 4C interacting domains compared to B cells (**Fig 5B**). The opposite is observed with the MiEκ 4C bait in B cells, where the enhancer is active and enrichment of ATAC-Seq peaks in 4C interacting domains is higher compared to T cells (**Fig 5B**). Thus, in line with previous studies using both HiC and 4C-Seq [15, 16], active regions of the genome preferentially contact other active regions while inactive regions contact other inactive regions, and this pattern is consistent across lineages.

**Fig 5:**
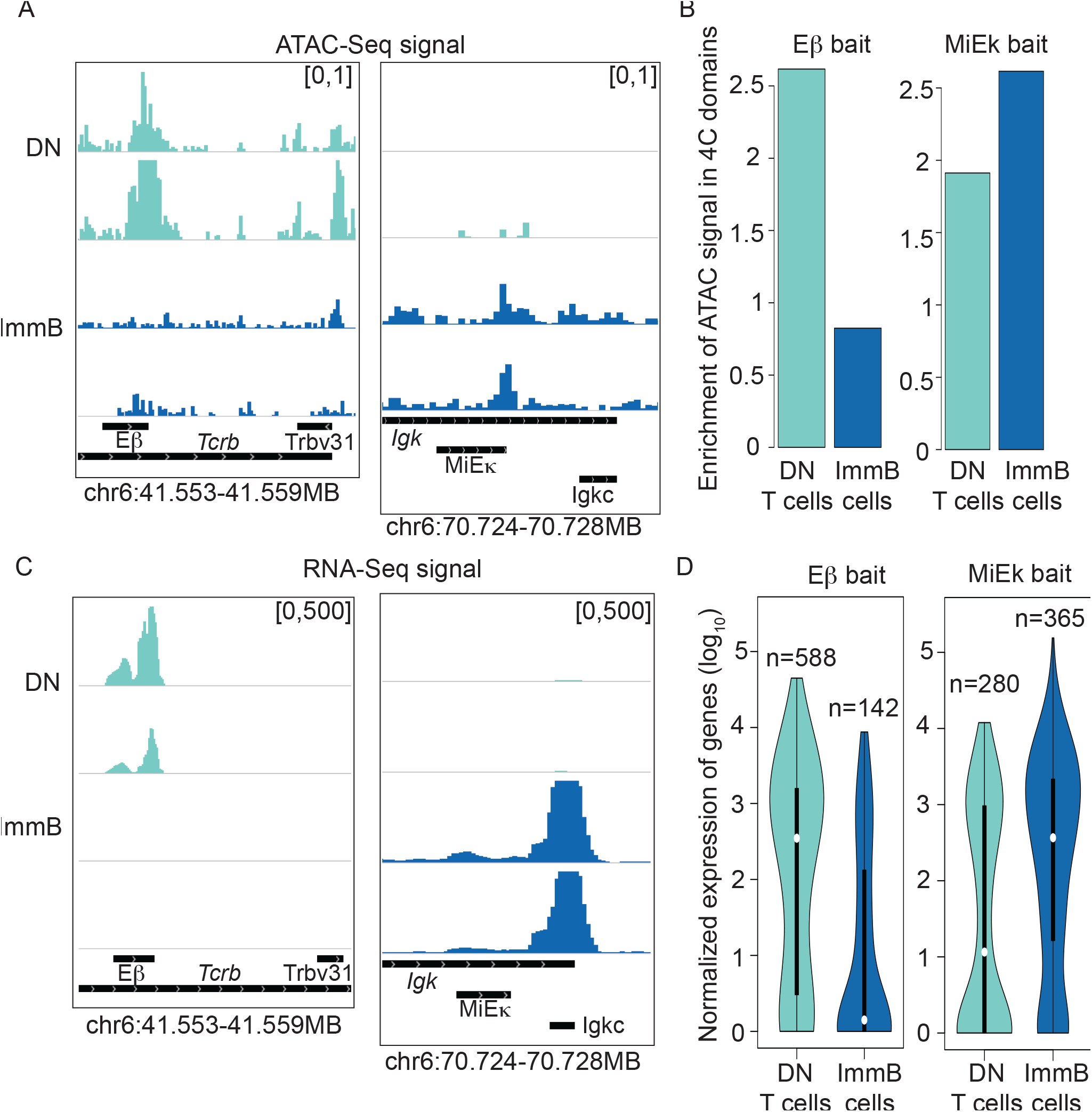
Regions with similar epigenetic and transcriptional status occupy the same nuclear space. (**A**) Normalized ATAC-Seq signal across the Eβ and MiEκ enhancer in T and B cells. (**B**) Enrichment of ATAC-Seq peaks in 4C interacting domains (**C**) Normalized RNA-Seq data for the *Tcrb* gene and the 3’ end of the *Igk* gene. (**D**). DESeq2 normalized expression values (log_10_) of genes that overlap with 4C-ker identified domains of interaction for each cell type.

To determine the relationship between transcriptional status and accessibility, we next integrated RNA-Seq data with the output from 4C-ker. We first confirmed the transcriptional activity of both enhancers across lineages, as demonstrated by the transcriptional activity of the*Tcrb* and *Igk* loci that are controlled by their respective enhancers (**Fig 5C**). The active Eβ enhancer selectively directs transcription of *Tcrb* in T cells, while MiEκ contributes to the high levels of *Igk* transcription that is found only in the B cell lineage. Next we compared the expression values of genes within the interacting domains across the different cell types. The genes within the Eβ-interacting domains in T cells show a higher transcriptional activity compared to genes within Eβ-interacting domains in B cells (**Fig 5D**). The reverse is observed in genes within MiEκ-interacting domains in B versus T cells (**Fig 5D**). Again, these results are in agreement with Hi-C studies, which show that regions with similar transcriptional activity occupy the same space in the nucleus [6, 15, 16].

## Discussion

Here we describe 4C-ker, a 4C-Seq analysis framework, that is unique in its ability to reproducibly detect short and long range-interactions on the same and across different chromosomes from a single viewpoint. Unlike other 4C-Seq pipelines, 4C-ker takes into account difference in coverage in regions proximal to the bait, far-*cis* and *trans*. As summarized in **Table 1**, 4C-ker outperforms all other methods in regions near the bait and in far-*cis* and performs comparably to the de Laat method for analysis of *trans* interactions. Moreover, all other tested methods fail to detect a region we defined as highly interacting using 4C-ker that we subsequently validated by FISH. Finally, 4C-ker also has the option to perform differential analysis of *cis* interactions.

**Table 1:** Summary of method comparison

|  | Near-bait | Far-*cis* | *Trans* | Differential analysis |
| --- | --- | --- | --- | --- |
| 4C-ker | Good | Good | Fair | Yes |
| **fourSig** | Majority of region called | Fair | Poor | No |
| **de Laat Method** | NA | Fair | Fair | No |
| **r3CSeq** | Restricted to the bait | Poor | Poor | Yes |
| **FourCSeq** | NA | NA | NA | Yes |
| **4cseqpipe** | Visualization | NA | NA | No |

4C-Seq can be used as an unbiased approach to identify short-range regulatory interactions that occur with the bait as well as long-range interactions that can provide insights into the global organization of chromatin in the nucleus. With 4C-ker, we validated long-range interactions from enhancer viewpoints and analyzed the epigenetic and transcriptional properties of interacting domains (where the validation includes reproducibility of results and experimental validation with FISH). This enabled us to demonstrate that the domains that 4C-ker calls have biological significance: active regions preferentially associate with active regions and inactive regions preferentially associate with inactive regions, as previously shown in Hi-C [6, 15, 16]. While Hi-C is limited in its ability to detect short-range interactions at low resolution, 4C-ker can identify both short and long-range interactions with higher resolution at lower sequencing depth.

One important consideration in 4C-Seq is to unravel how the profile of interactions generated in a population of cells relates to the physical constraints of chromosomes within the nucleus. For example, we need to better understand the implications of the differences in 4C-Seq profiles when an active or an inactive bait is used. Reduced interactions from an inactive bait likely reflect a less mobile compacted chromatin structure that could be embedded within the chromosome territory. To explore these relationships we need improved pipelines for integrating other genome wide techniques such as RNA-Seq, ATAC-Seq, and ChIP-Seq with 3C-based data sets. Only then can we learn whether inactive regions of the genome interact with regions that share epigenetic modifications and are bound by common regulatory factors as has been shown for active regions that are co-regulated [26, 27].

Although 4C-Seq only provides information on interactions from a single viewpoint, it can help to identify intricate loop structures at a finer resolution than Hi-C, and this in turn will provide a basis for understanding regulatory interactions. Furthermore, it can identify long-range interactions in *cis* and in *trans* that likely reflect inter-TAD interactions on the same or different chromosomes. These interactions need to be validated by FISH analysis, which in contrast to chromosome conformation capture, faithfully reflects the appropriate chromatin compaction state and recapitulates the findings from individual live cells (as opposed to averaging over populations) [28]. Furthermore, FISH analysis can provide information about whether a particular region is embedded within a chromosome territory or looped away, which can be reflective of gene activity or association with repressive pericentromeric heterochromatin [29].

The 4C-ker pipeline can be adapted for analysis of data from new 3C-based techniques such as Capture-C [30], T2C [31] and CHi-C [32], that use oligonucleotides to enrich interacting fragments from multiple baits in a single experiment. Furthermore, the high resolution of 4C-Seq data can be used for determining the finer structure of domains identified with Hi-C. Finally, it should be pointed out that there is a great deal of variability between 4C-Seq experiments generated by different labs, and it is clear that the field would benefit from standardized protocols and quality control of datasets that lend themselves to comparisons between experiments from different sources. Going forward 4C-ker will provide a much-needed tool for comprehensive analysis of 4C datasets derived from different experimental approaches.

## Material and Methods

### Ethics Statement

Animal care was approved by Institutional Animal Care and Use Committee. Protocols number is 150606-01 (NYU School of Medicine). The authors have no conflict of interest.

### Mapping 4C-Seq reads to a reduced genome

The sequence reads generated from a 4C-Seq experiment typically contain the primer sequence ending in the primary restriction enzyme followed by the interacting fragment captured by the bait. The portion of the read following the restriction enzyme sequence is mapped to a reduced genome — a set of unique sequences (with the same length as the interacting fragment sequenced) that are directly adjacent to all sites in the genome of the primary restriction enzyme used. We define these unique sequences as ‘potential fragments.’ We used oligoMatch (from UCSC command line tools) to find all the primary restriction enzyme recognition sequences in the genome and a custom shell script (provided) was used to create the reduced genome. Reads were mapped to the reduced genome using Bowtie2 [32] (command-line options: -N=0, in addition -5 was used to trim the barcode and primer sequence).

We define the fragments in the reduced genome that have at least 1 read mapped to it as an ‘observed fragment’. The read count at each observed fragment is extracted from the Bowtie output (SAM file) and transformed to a bedGraph file (4 columns with chr,start,end,count at each observed fragment) that can be uploaded to IGV for visualization. A custom shell script is provided to generate these bedGraph files. For paired-end sequencing experiments the read containing the bait and the primary restriction enzyme was mapped as single-end data.

### Dynamic window sizes to correct for coverage

Adaptive window sizes were determined using the k-th nearest neighbor approach to account for the change in 4C-Seq coverage in different regions. The value of k determines the number of observed fragments to be analyzed within each window. The window size is determined for each observed fragment as the linear distance to the k-th nearest observed fragment, which will result in a larger window size in regions where few fragments are observed and vice versa. On the bait chromosome, the adaptive windows are determined left and the right of the bait starting from the bait coordinate. For example, the first window on the left of the bait will be based on the k observed fragments on the left of the bait and the first window on the right will be based on the k observed fragments on the right of the bait. In this manner, the windows are built until the end of the chromosome. For *trans* interactions, this process is performed independently for each chromosome starting with the first fragment identified at the beginning of the chromosome. Window sizes are determined for each sample in a given dataset. Then a smooth spline (smooth.spline function in R with a smoothing parameter of 0.75) is fitted to the window sizes separately for each chromosome in order to get a window size at each position along the chromosome that can be used for the entire dataset (**S1A Fig**).

To build the final windows we use overlapping windows to more accurately identify the borders of interacting domains. ***Cis:*** Using the bait coordinate, the size of the first window is predicted from the fitted spline and this is used to build a window to the left and right of the bait. Adjacent windows start at the mid-point of the bait window and the size is again determined by the fitted spline. In this manner, overlapping windows are generated for the region near the bait or the entire chromosome and will be used to analyze interactions for all samples in the given experiment. For the analysis near the bait we used k=5 and, when analyzing the entire bait chromosome, k=10. ***Trans:*** Starting from the beginning of each *trans* chromosome, we predict the window size from the fitted spline. The next window starts from the mid-point of the first window and this process continues to the end of the chromosome. We used k= 15 for all *trans* analysis using 6bp cutters and k=100 for 4bp cutters.

### Log_10_ transformation of normalized counts within windows and distance from the bait

To reduce the effect of PCR artifacts, fragments with counts greater than the 75^th^ quantile within a given window are trimmed to this value. In this manner, we reduce the value of single fragments that have an extremely high count that are not supported by neighboring fragments with a similar signal. The counts at observed fragments within each window are normalized across all samples in the dataset using the method described in the DESeq2 [17] R package where each window is considered as a feature (or gene). For windows in *cis*, the distance from the bait to the mid-point of each window (in bp) is also calculated. A pseudo-count of 1 is added to the normalized window counts and the distance value followed by a log_10_ transformation of the values. The log-transformation of the data results in an approximately linear function that describes the decrease in counts as the distance from the bait increases.

### Hidden Markov Model

In order for 4C-ker to take into account conditional dependencies among neighboring genomic elements we propose to use a three-state Hidden Markov Model (HMM) where the hidden states represent genomic regions that show high frequency of interactions in the population (high interaction region-HI), low frequency (low interaction region-LI), and no significant frequency of interactions (no interaction-NI) with the bait. A separate model was learned for *cis* and *trans* chromosomes. We used the depmixS4 R package [33] to specify and train the described HMM.

#### Near-bait and cis Parameters

For *cis* interactions we propose a covariate-adjusted HMM. We denote the number of windows on the *cis* chromosome as T. The input data consists of the observed log-normalized counts **O**_1:*T*_ = {*O*_1;_…, *O_t_*,…, *O*_T_} where *O_t_* = 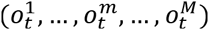, denotes the counts for window t from M biological replicates. The hidden states are denoted by *S*_1:*T*_ = {*s*_1;_ *s*_2_,…, *s_t_*,…, *s_T_*}. We use the log_10_-transformed distances from the mid-point of each window to the bait *D*_1:*T*_ = {*d*_1_ *d*_2_,…, *d_t_*,…, *d_T_*} as covariates. For the *cis* interaction three-state HMM the joint likelihood of observations and hidden states, given model parameters *θ* and covariates *D*, is

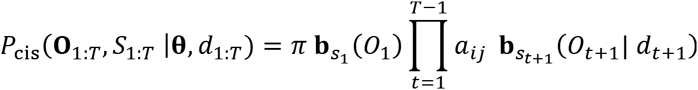

with the following components:

1. Hidden states *s_t_ ∈* {1,2,3} (1=no interaction, 2=low interaction, 3=high interaction).
2. The initial state distribution **π** with elements *π_i_* = *P[*s*_t_* = *i*], 1 ≤ *i* ≤ 3.
3. The state transition matrix *A* = {*a_ij_*} with unknown entries *a_ij_* = *P*[*s_t+1_* = j| *s_t_* = *i*], 1 ≤ *i, j* ≤ 3.
4. Emission probabilities (observation densities) are represented as vector **b***_S_t__* with elements 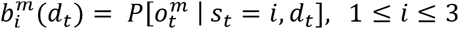 that model the conditional density of observations 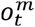 in window t in the m^th^ replicate.

We use the following linear model with a Gaussian response function to link count data and distance covariates *D_1:T_*

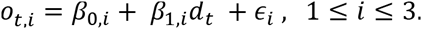

We assume that counts *o_t,i_* in state i are normally distributed with unknown mean and variance *σ_l_^2^*

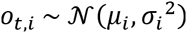

The expected value of counts *μ_i_* is thus a linear function of the distances, controlled by the parameters **β**_i_ = (*β_0, i_, β_1, i_*).

The resulting *cis* model comprises a total of 21 parameters **θ** = {***π**, A*, **β**_1_, **β**_2_, **β**_3_, *σ*_1_, *σ*_2_, *σ*_3_}.

#### Synthetic training data and parameter estimation

**Training data:** 4C-Seq signal can vary based on the activity of the bait, location on the chromosome, and possibly the species in which the experiment is being done. Therefore a different set of parameters is learned for each dataset. To simulate the unknown underlying population of 4C-Seq data we create input data Õ_1:*T*_ for the *cis* model by generating bootstrap samples from the biological replicates. For each non-overlapping window along the *cis* chromosome we randomly draw a 4C-Seq signal from the m replicates. The synthetic samples then undergo the same normalization procedure as the original replicates to get the counts per window and the linear distance to the bait. This method of generating synthetic samples allows us to generate training data that have transitions different from the observed data (biological replicates).

***Consistency constraints:*** The overwhelming signal in the bait region can sometimes lead to unsuitable model parameter estimates that do not describe the three states correctly. To achieve state-consistent estimates of decrease in signal with increasing distance from the bait, we imposed the following set of linear constraints on the emission parameters:

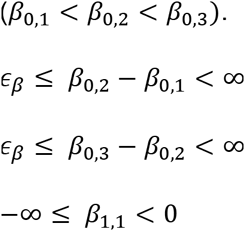

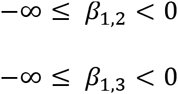

These constraints ensure that (i) expected emission probabilities strictly decrease with distance from the bait and (ii) the non-interaction, low-interaction, and high-interaction states obey the correct ordering. We set the value of the slack variable *∊_β_* = 0.1.

**Initial parameter values θ_0_**: In order to find reasonable starting values for the parameters **θ**, we performed a parameter sensitivity analysis by fitting the HMM to the *CD83* HindIII datatset using 1000 random starting parameter values and determining the parameter region that resulted in reproducible results (**S5 Fig**). This analysis resulted in using 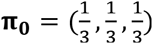 and initial transition probabilities *a_ii_* = 0.5 , *a_ij_* = 0.25, *i ≠ j* (**Fig 1**). For the emission probabilities, we divide the chromosome into 30-window segments and separate the counts in each segment by those lower than the 60^th^ quantile (no interaction), those between the 60^th^ and 90^th^ quantile (low-interaction) and those greater than the 90^th^ quantile (high-interaction). These counts are then used to estimate the starting values for the emission probabilities. In order to ensure that the parameters are not close to the boundaries of the constraints, we set *β*_0,2_ = 0.8 * *β*_0,3_ and *β*_0,1_ = 0.5 * *β*_0,3_. The predicted counts from the estimated linear model for each state along the *cis* chromosome are plotted in **S6A Fig**.

**Maximum-likelihood estimation:** A general nonlinear augmented Lagrange multiplier method solver (solnp function in R package Rsolnp) was used to find the maximum likelihood estimates of the parameters with the imposed linear constraints. The average estimated initial state and transition probabilities are shown in the HMM model in **Fig 1**.

#### Defining domains that are in close proximity to the 4C-Seq viewpoint

After model inference we use the Viterbi algorithm to assign the interaction states to each of the windows on the biological replicates. If adjacent overlapping windows are assigned to different states, we trim the window called as high-interaction region in order to retain the part of the window not in a conflicting region (**S6B Fig**). Overlapping windows called as highly interacting are merged to define large domains of interaction with the bait. The final set of highly interacting domains for a given 4C-Seq data set is the intersection of the trimmed windows across all replicates.

#### Trans

We learned one model for all *trans* chromosomes where the input to the HMM is the normalized counts for each window. Let **O**_1:*T_N_*_ 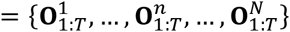 represent the counts across all windows and replicates in all N *trans* chromosomes. For the *trans* model we used a three-state HMM *without* covariate adjustment for distance. The joint likelihood of observations and hidden states, given model parameters *θ* thus reads

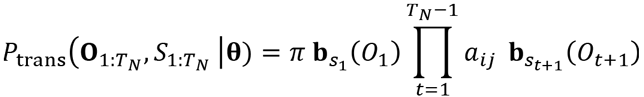

We again assume the normalized counts to be multivariate normal. The resulting *trans* model without covariate adjustment thus comprises 18 parameters **θ** = {**π**, *A*, μ_t_, μ_2_, μ_3_, σ_x_, σ_2_, σ_3_}.

#### Synthetic training data and parameter estimation

A synthetic sample was built such that each chromosome was randomly selected from the pool of replicates. The following constraints were added to ensure that the parameter values are suitable to distinguish between the states.

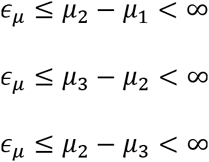

We set the value of the slack variable *∊_μ_* = 0.05 and used the depmixS4 R package to specify and train the described HMM where again the general nonlinear augmented Lagrange multiplier method solver solnp was used to find the maximum likelihood estimates of the parameters with the imposed linear constraints.

### Quantitative analysis using DESeq2

For multiple conditions with the same bait, a merged set of domains is generated that contains those called as highly interacting in at least one of the conditions - Dataset-specific interacting domains (DIDs). We can then obtain the raw window counts with each domain and use DESeq2 to perform a quantitative differential analysis. DESeq2 has been developed primarily to analyze RNA-Seq data but can also be applied to any count dataset that follows a negative binomial distribution. Therefore, we decided to use the method to look for quantitative differences between conditions in 4C-Seq.

### Similarity index

Similarity index was calculated based on a previously described method for dealing with more than 2 replicates [34].

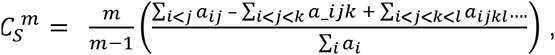

where *m* is the number of replicates in the dataset and *a_ij_* is the sum of the size of overlapping HI domains between replicate *i* and *j* and £*_i_a_i_* is the sum of the size of the merges HI domains from all replicates. Domains from both replicates were retained when 50% of the domains overlapped with the other replicates. When comparing with different number of replicates, we divide by m to get a score between 0-1.

### Comparison to other methods

The interactions defined for each replicate by the four methods were used to calculate the similarity index.

**fourSig**: 4C-Seq data was mapped to mm9 genome. A window size of 5 was used for the analysis near the bait, and a window size of 31 was used for far-*cis* and *trans* analysis with 1000 iterations and an FDR cut-off of 0.05 (liftOver was used to convert the results to the mm 10 genome).

**The de Laat method**: The input for this method was generated using our alignment pipeline. Domains were called based on the significant contacts.r file.

**r3CSeq**: The program currently does not allow for analysis with the mm10 genome, therefore we mapped the 4C-Seq data to the mm9 genome. Since the workflow for “working with replicates” requires a control and condition experiment, each replicate was run through the “work without replicates” pipeline. The data was analyzed at the level of restriction fragments, 20Kb windows and 100kb windows. Only the results from the fragments analysis are shown as this was deemed to be the most optimal for high resolution and had more interactions called in far-*cis* and *trans* (liftOver was used to convert the results to the mm10 genome).

**FourCSeq**: 4C-Seq data was mapped to the mm10 genome. No significant interactions were observed with an FDR cut off of 0.1. Although we were not able to identify any interactions with the datasets used in this study, we were able to reproduce their results with the example dataset provided.

### 4C-Seq experiments

Details of publically available datasets downloaded for this study can be found in **S1 Table**. We used datasets that had more than one replicate available in GEO and processed the FASTQ files using our pipeline. Datasets were further excluded if less than one million reads were available after removal of undigested and self-ligated 4C fragments. Samples were also required to have at least 40% of the reads on the *cis* chromosome and 40 % coverage in the 2Mb region around the bait for 6bp cutters and 200kb for 4bp cutters as this is considered a standard quality control for a good 4C experiment [35]. Basic statistics for the datasets used can be found in **S1 Table**. All datasets generated for this study can be found in GEO (GSE77645).

The following datasets were generated from mouse cells for this study. *Cd83, Igh-*Cγ1 baits on activated mature B cells, *Igk* MiEκ, *Tcr* Eβ bait in double negative (DN) T cells and immature B cells. See **S1 Table** for details of primers and enzymes used for these experiments.

The 4C-Seq protocol was performed as described previously [14] and libraries were sequenced using the HiSeq2500 Illumina platform. Splenic mature B cells were isolated and induced to undergo class switch recombination as previously described [14]. Cells were collected on day 2 of activation. DN T cells, and immature B cells were isolated as described before [29, 36] and pooled to obtain 10 million cells for each replicate at each developmental stage.

### FISH validation

Activated mature B cells for FISH analysis were isolated as described above. 3D-FISH was performed as described previously [37]. Interphase cells were analyzed by confocal microscopy on a Leica SP5 AOBS system (Acousto-Optical Beam Splitter). Optical sections separated by 0.3μm were collected using Leica software and only cells with signals from both alleles (>95% of cells) were analyzed. Separation of alleles was measured in 3D from the center of mass of each signal using Image J software.

### ATAC-Seq

DN T cells as well as immature B cell were isolated as described above. ATAC-Seq was performed in duplicate as described previously [38] with the following modifications: libraries were amplified with KAPA HiFi polymerase. Libraries were sequenced with HiSeq using 50 cycles paired-end mode. 50bp-paired-end reads were mapped to mm9 using Bowtie2 with the following parameters: –maxins 2000, –very-sensitive Reads with MAPQ score < 30 were filtered out with Samtools, and duplicate reads were discarded using Picard tools. For each sample condition, biological replicates were merged with Samtools, and peaks were called using Peakdeck [39] with the following parameters: -bin 75, -STEP 25, -back 10000, -npBack 100000. Peaks were further filtered to a raw p-value cutoff of 1E-4 (liftOver was used to convert the results to the mm10 genome). A custom script was used to determine peak maxima, and maxima were extended by 50bp on either side to yield peaks of ∼100bp.

### RNA-Seq

DN and immature B cell were isolated as described above. RNA-seq libraries were prepared as previously described using the Ribo-Zero kit for depletion of ribosomal RNA [40]. Reads were mapped using Tophat version 2.0.6 [41]with the following parameters: –no-coverage-search -p 12 –no-discordant –no-mixed -N 1 –b2-very-sensitive. Number of reads per gene (RefSeq annotation) was calculated using HTSeq-count [42]. Normalization of counts per gene was done using DESeq2.

## Acknowledgements

We would like to thank members of the Skok and Bonneau lab for helpful suggestions and discussions. We would like to thank Lili Blumenberg for comments on the manuscript. We would also like to thank Shenglong Wang from the NYU HPC facility for timely technical support, NYU CHIBI and the NYUMC sorting and genome facilities. This work was supported by grants from the NIH (GM086852 (JS) and GM112192 (JS and RB). GM32877-21/22, PN2-EY016586, IU54CA143907-01 and EY016586-06 (RB). and NSF IOS-1126971 (RB). JS is a Leukemia Lymphoma Society Scholar, PR is a National Cancer Center postdoctoral fellow and an American Society of Hematology Fellow.

## Author contributions

RR developed and implemented 4C-ker. RR, PR, CM, EM, RB and JS designed the concept of the study. RR, PR, CM, EM and SB analyzed the data. RR, PR, YF, ES, CP and VS generated FISH, ATAC-Seq and 4C-Seq data. RR, PR,CM and JS wrote the manuscript with comments from all authors.

---

**Fig S1.**
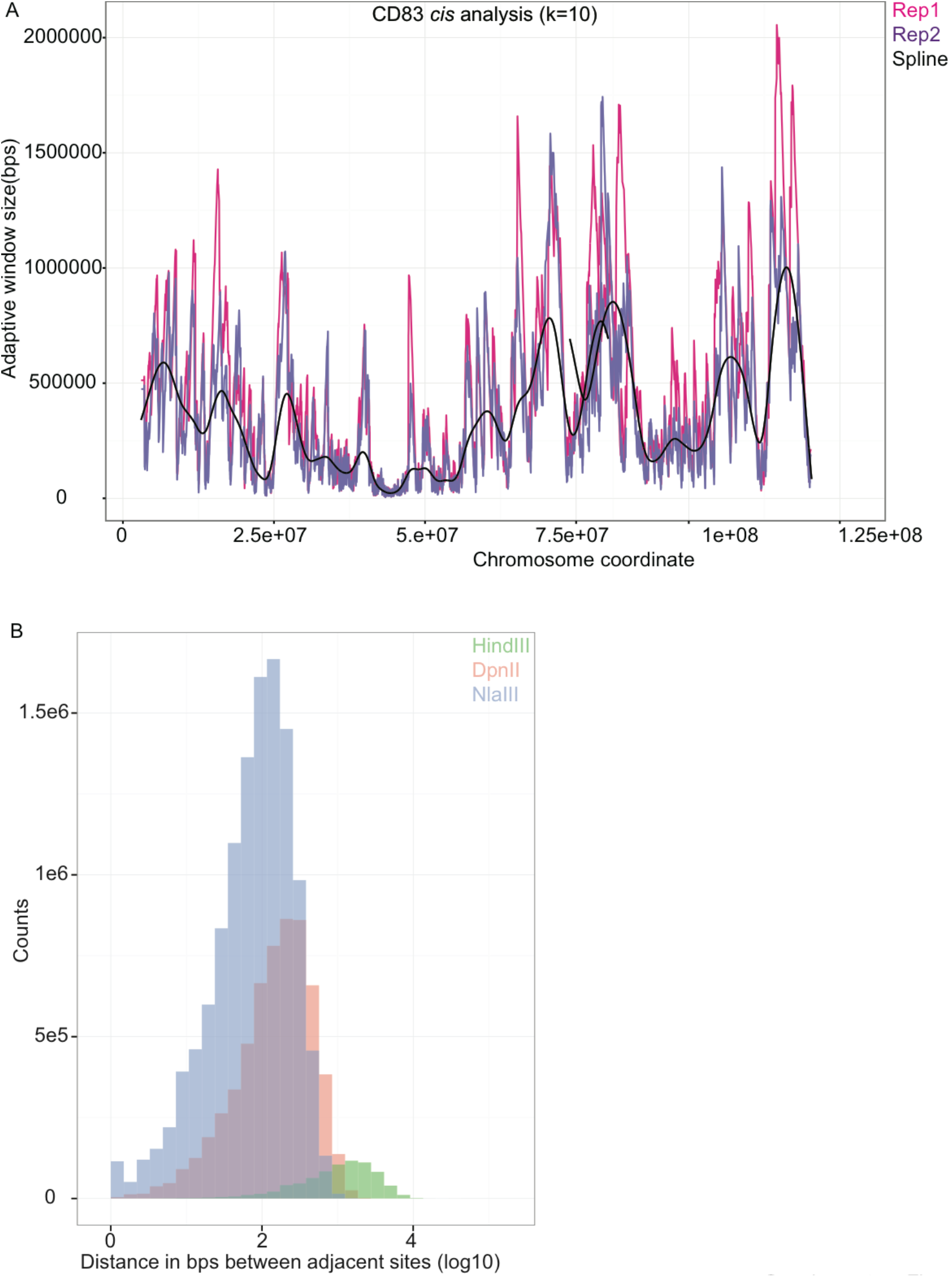
**(A)** Adaptive window analysis for the *cis* chromosome using k=10. A spline is fitted to the two replicates (*Cd83* dataset) to generate a smooth curve which is then used to create the overlapping windows. **(B)** Histogram of the distance between adjacent restriction enzyme sites in the mouse mm10 genome. NlaIII has the highest number of sites in the genome resulting in shorter distances between adjacent sites.

**Fig S2.**
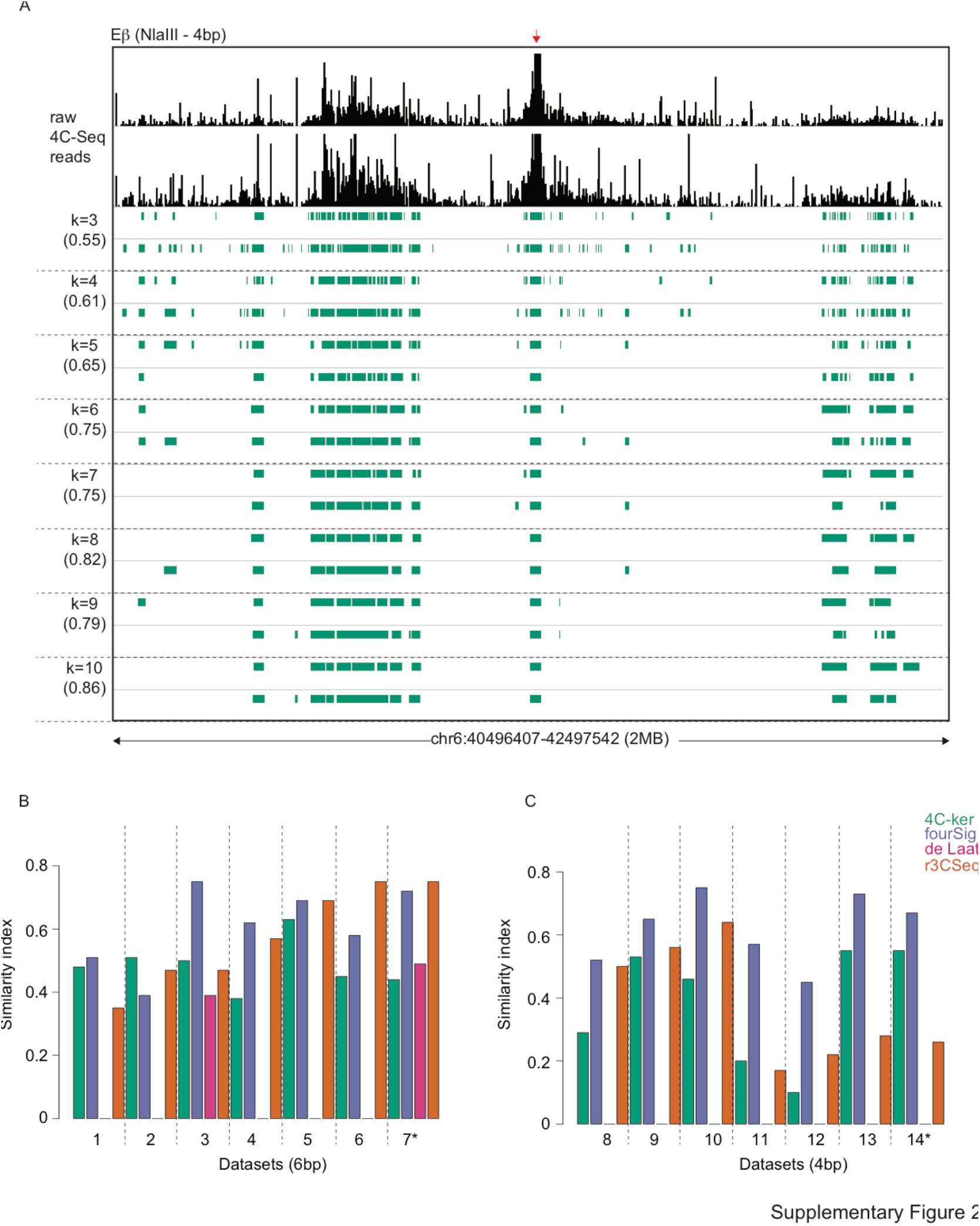
(**A**) Near bait analysis using different values of k for the 2MB region around the Eβ bait in DN cells. Domains called for the two replicates are shown and the similarity index below each value of k. (**B-C**) Similarity index between replicates for interacting domains identified in the region around the bait for 6bp and 4bp cutter datasets respectively. The numbers in the x-axis refer to each dataset. * Represents datasets shown in **Fig 2A**.

**Fig S3.**
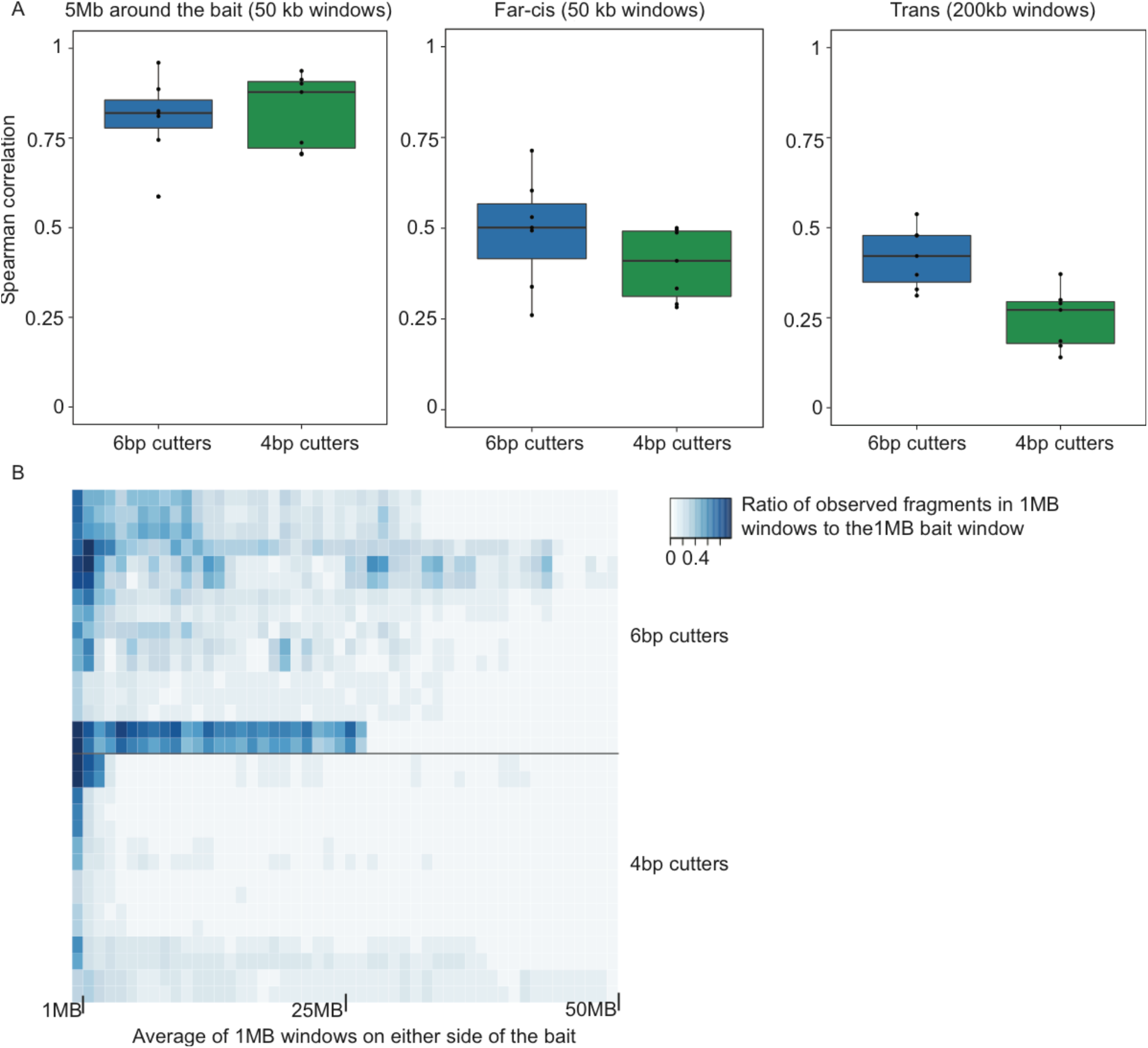
**(A)** Raw counts for different window sizes were used to calculate Spearman correlation across several datasets (listed in Supplementary Table1). The mean of pairwise correlations were plotted for datasets with greater than 2 replicates. **(B)** Ratio of observed fragments in 1 MB windows (up to 50MB away from the bait) against the observed fragments in the 1MB window encompassing the bait. Each row represents a 4C-Seq experiment (replicates are separated).

**Fig S4.**
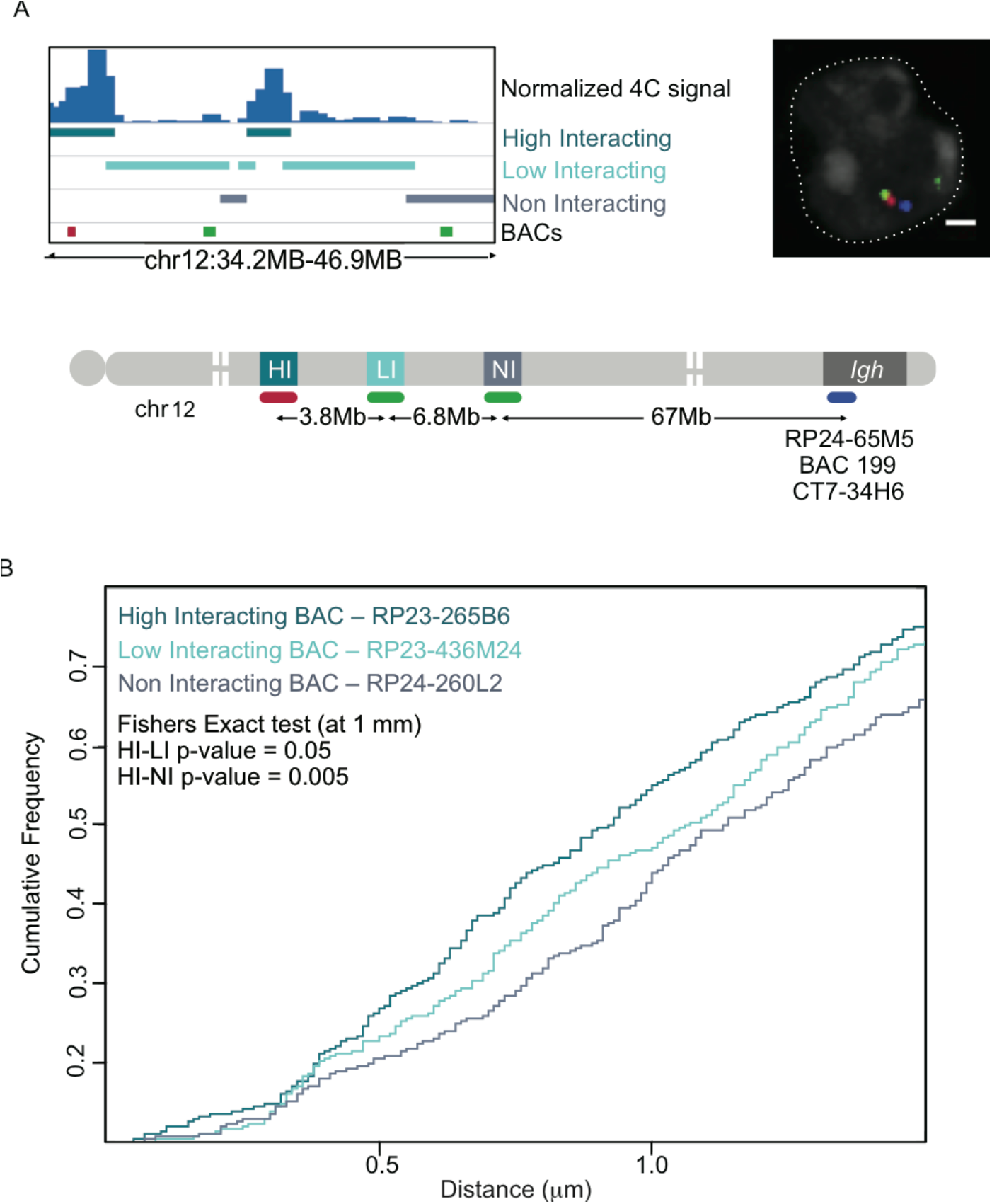
(**A**) Browser view of a far-*cis* region on chromosome 12 showing domains identified as High, Low and Non interacting states and the location of BACs chosen to label these regions as well as the distances separating them from each other and from *Igh*. These BACs, together with probes labeling the constant region of *Igh* were used for 3D-FISH on activated B cells. (**B**) The distance from each BACs to *Igh* was measured and plotted as a cumulative frequency curve. A shift to the left represents closer proximity to *Igh*. The BAC representing the High interacting state is more frequently found closer to *Igh* than the BACs representing the Low and Non interacting states. This difference is statistically significant using a Fisher’s exact test at 1μm distance. The FISH example shows one Z plane where one chromosome 12 is visible.

**Supplementary Fig 5:**
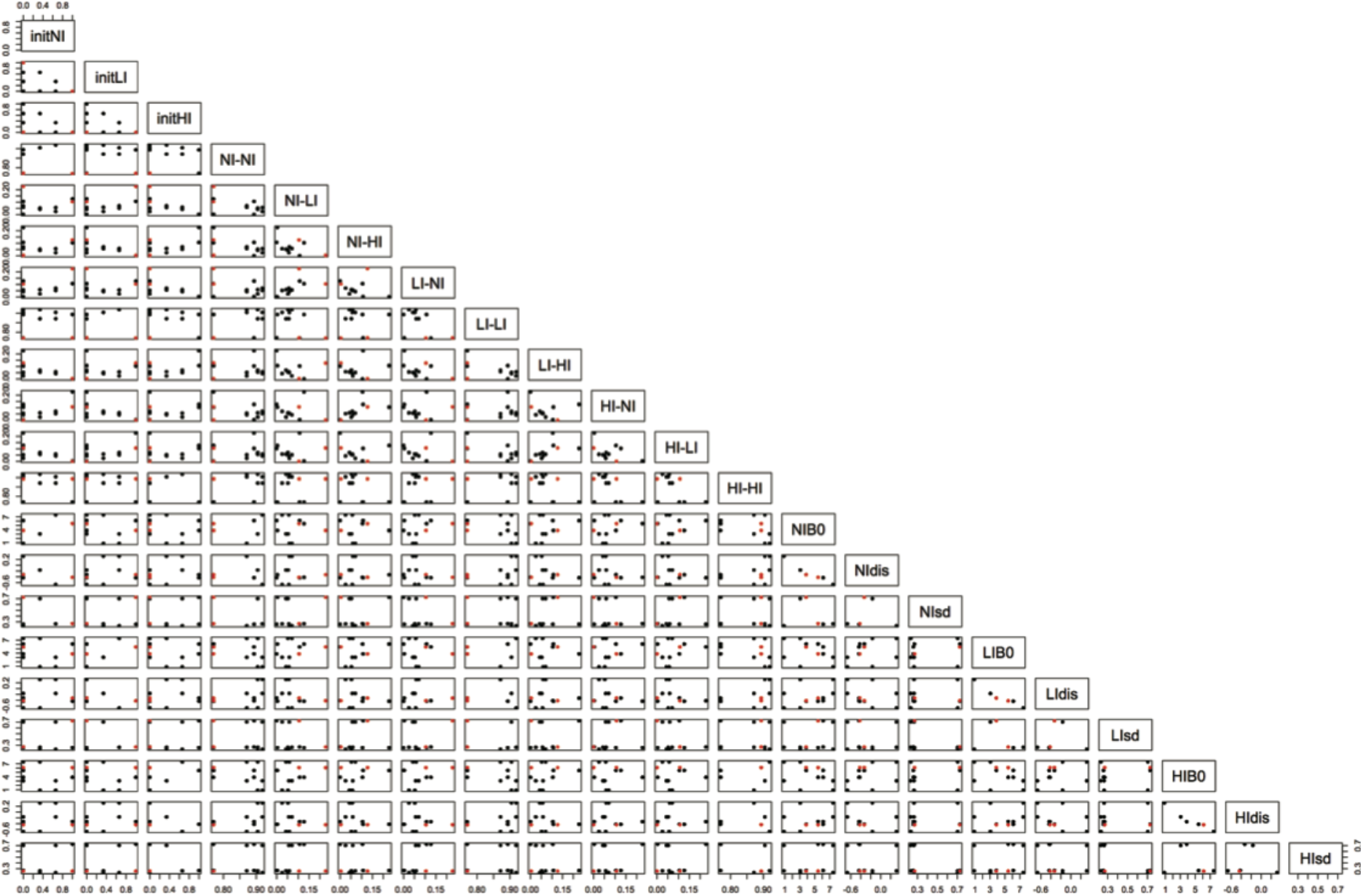
Results of parameter estimates using 1000 different starting values. Estimation was performed using the EM algorithm with no constraints. The set of parameters that resulted in Viterbi calls with a reproducibility of 60% or greater across replicates are colored in red. The probability of transitioning to the same state is always higher than transitioning to a different state. As expected, the distance covariate term (names here as dis) is always negative for the reproducible set of parameters, confirming the decrease in signal with increase linear distance from the bait.

**Supplementary Fig 6:**
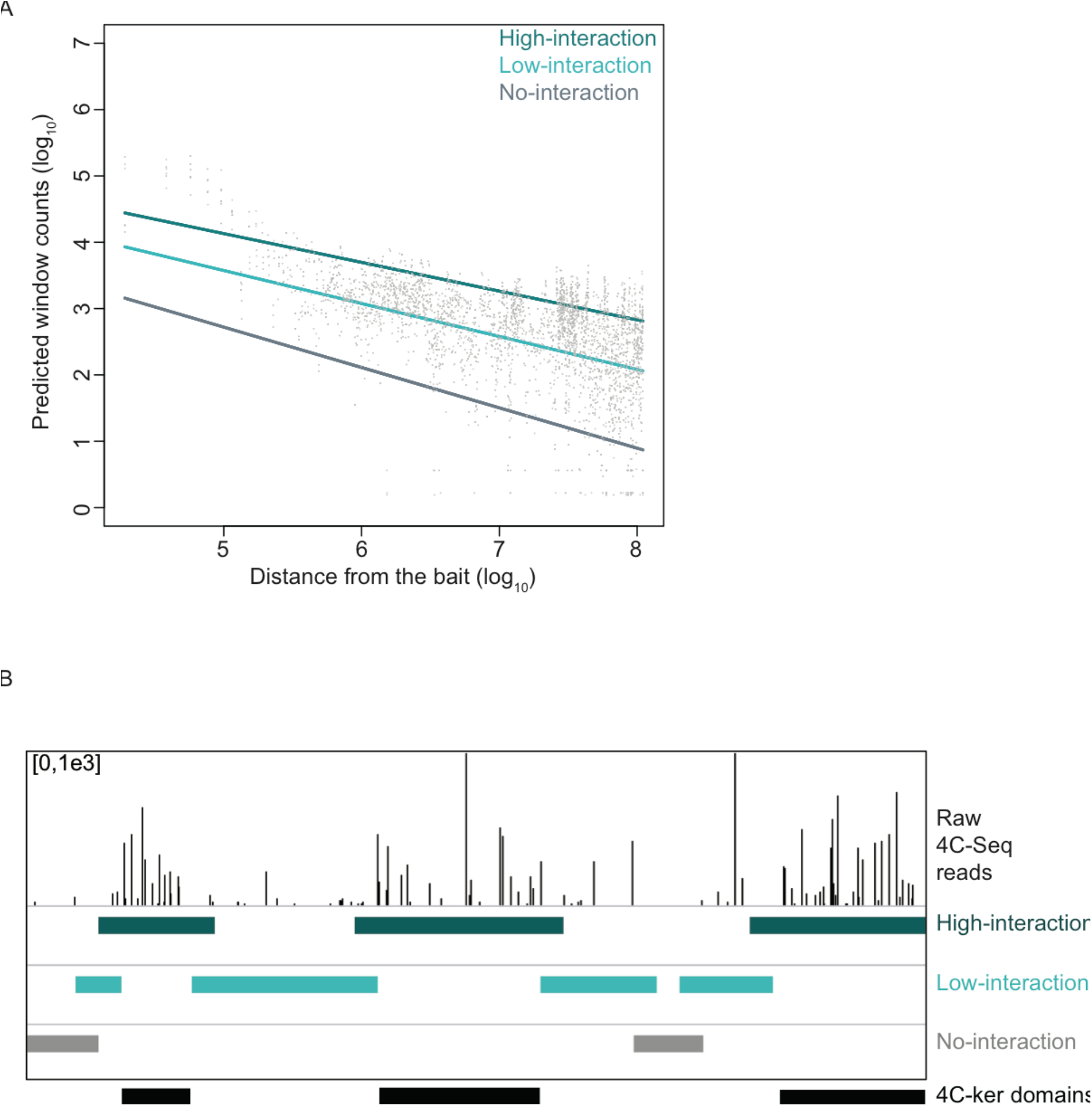
Results of the HMM. (**A**) Using the distance from the bait, the window counts were predicted from the estimated linear model for each of the HMM states. (**B**) Region of the bait chromosome showing the hidden states inferred by the Viterbi algorithm and the trimmed 4C-ker domains.

**Table S1.**
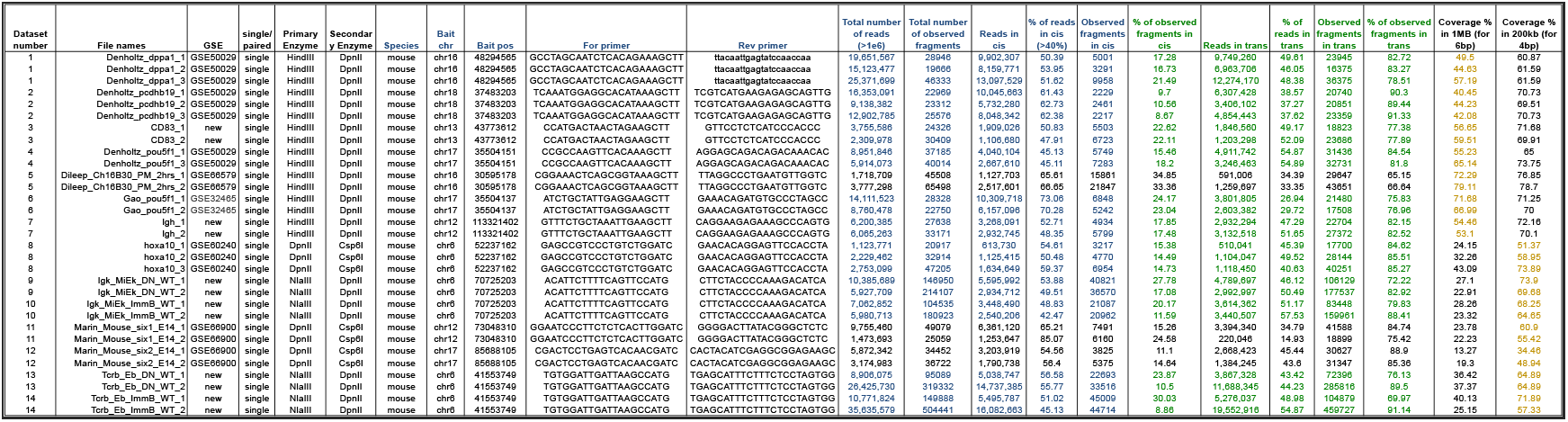
Description of datasets used for this study.

